# Somatic small-variant calling methods in Illumina DRAGEN™ Secondary Analysis

**DOI:** 10.1101/2023.03.23.534011

**Authors:** Konrad Scheffler, Severine Catreux, Taylor O’Connell, Heejoon Jo, Varun Jain, Theo Heyns, Jeffrey Yuan, Lisa Murray, James Han, Rami Mehio

## Abstract

We present the DRAGEN™ somatic pipeline for calling small somatic variants from tumor samples, with or without paired normal samples. The DRAGEN somatic variant caller offers 1) a flexible architecture that can be used on a wide array of somatic use cases; 2) built-in noise models enabling robustness against various sources of noise artifacts (mapping, genome context, or sample specific); 3) performance of joint analysis of tumor and normal samples in the case of a tumor-normal workflow yielding improved accuracy; 4) benefits from FPGA acceleration for efficient run time. We demonstrate the speed and accuracy of the DRAGEN tumor-normal pipeline across a range of whole genome sequencing (WGS) datasets and compare against third party tools such as Mutect2/GATK4 [1] and Strelka2 [2]. DRAGEN secondary analysis outperforms all other tools with its ability to complete a 110x/40x T/N whole-genome analysis in less than two hours. It offers exceptional accuracy, with higher sensitivity and precision than third party tools. We also show that the DRAGEN T/N workflow supports analysis of liquid and late-stage solid tumors by tolerating tumor-in-normal (TiN) contamination.

## 1 Introduction

Somatic mutations (single nucleotide variants (SNVs) and indels (insertions and deletions)) are frequently found in cancer genomes [3]. Large-scale analysis from The Cancer Genome Atlas (TCGA) has revealed that there are about 125K and 8K unique somatic SNVs and indels, respectively, found across 7,000 cases, for the 10 most common tumor types (including lung, breast, brain, and colorectal) [4]. SNVs and/or indels in genes such as BRCA1, BRCA2, and EGFR exon 20 involved in either DNA damage repair or activation of oncogenic pathways are well documented and serve as biomarkers for therapeutic interventions.

Somatic mutation detection plays a crucial role in improving our understanding of the disease and patient outcomes. It drives cancer research by identifying specific genetic changes in cancer cells. This information helps in developing new tests, treatments, and prevention strategies tailored to the mutations, as well as stratifying cancer patients for precise and effective treatment.

The field of somatic variant calling is diverse, encompassing a range of sequencing approaches. Panel-based tumor-only sequencing focuses on a limited set of cancer-related genes and is used to identify mutations in these genes. On the other hand, whole exome sequencing (WES) and WGS, with or without a matched normal sample, provide a more comprehensive view of the genomic changes in cancer cells. Each of these assays has its own set of requirements, including sample quality, sequencing depth, and bioinformatics analysis pipelines, that need to be met to accurately identify and interpret the somatic mutations. The choice of assay depends on various factors, such as the research question, type of cancer, and available resources.

These oncology applications have stringent requirements in terms of sensitivity and precision. For example, the Illumina TruSight Oncology (TSO) 500 panel for solid tumors has a limit of detection (LOD) of 5% variant allele frequency (VAF), with analytical sensitivity and specificity of 96% and 99.9995%, respectively [5]. The TSO500 ctDNA assay has a LOD of 0.5% VAF, with analytical sensitivity and specificity of 95% and 95%, respectively [6]. With high tumor content (> 50%), the LOD of WGS T/N ~100x/40x is > 5% VAF and with WES T/N it can go as low as 1 to 2% VAF with high read coverage (>100%) [7].

DRAGEN secondary analysis is a unified solution that addresses the various requirements of different somatic variant calling assays. It was designed to streamline the process and equip researchers with a common pipeline, eliminating the need for multiple solutions. Equipped with advanced noise models, DRAGEN delivers a more sensitive and precise method for identifying and characterizing cancer-related mutations. We describe those methods below and show benchmarking accuracy and run time results.

## 2 DRAGEN Somatic Pipeline

The DRAGEN somatic pipeline identifies somatic variants that can exist at low allele frequencies in tumor samples. The pipeline can analyze tumor/normal pairs (Fig. 1A) or tumor-only sequencing data (Fig. 1B). The somatic pipeline makes no ploidy assumptions about the tumor sample, allowing sensitive detection of low-frequency alleles.

**Figure 1.**
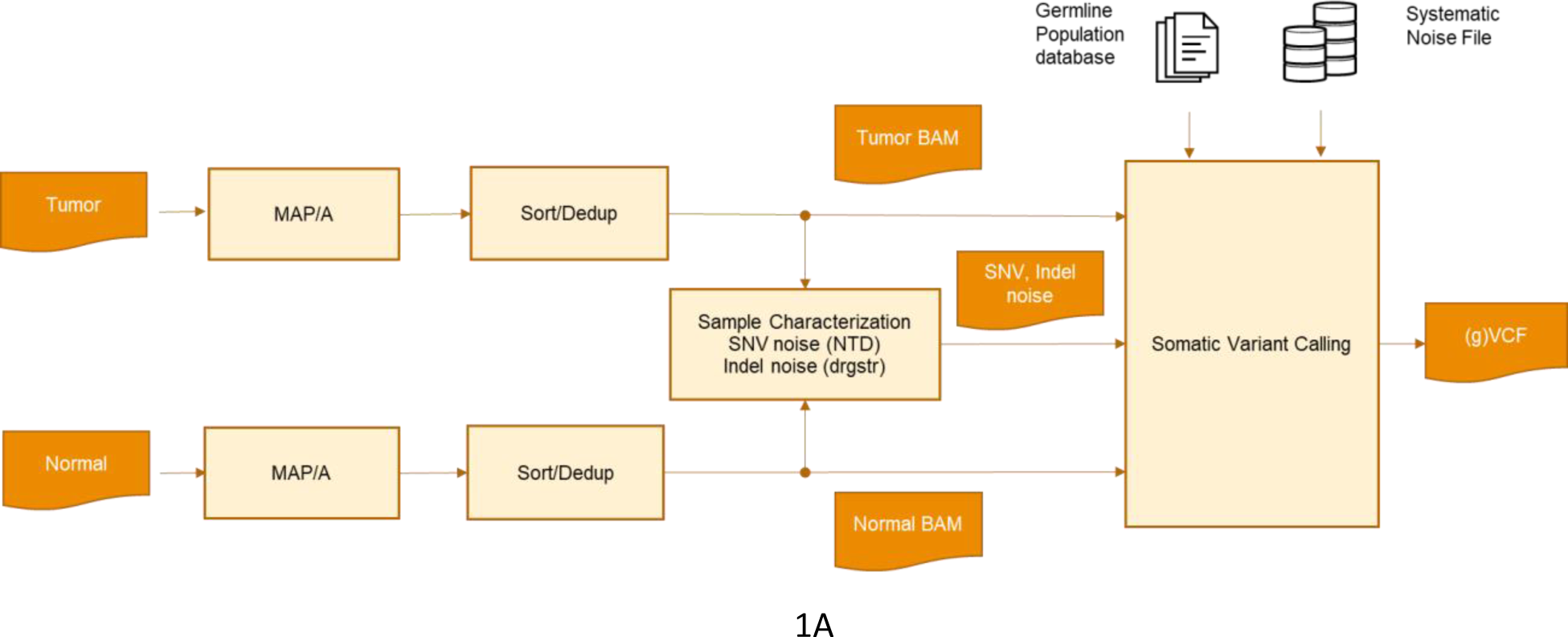
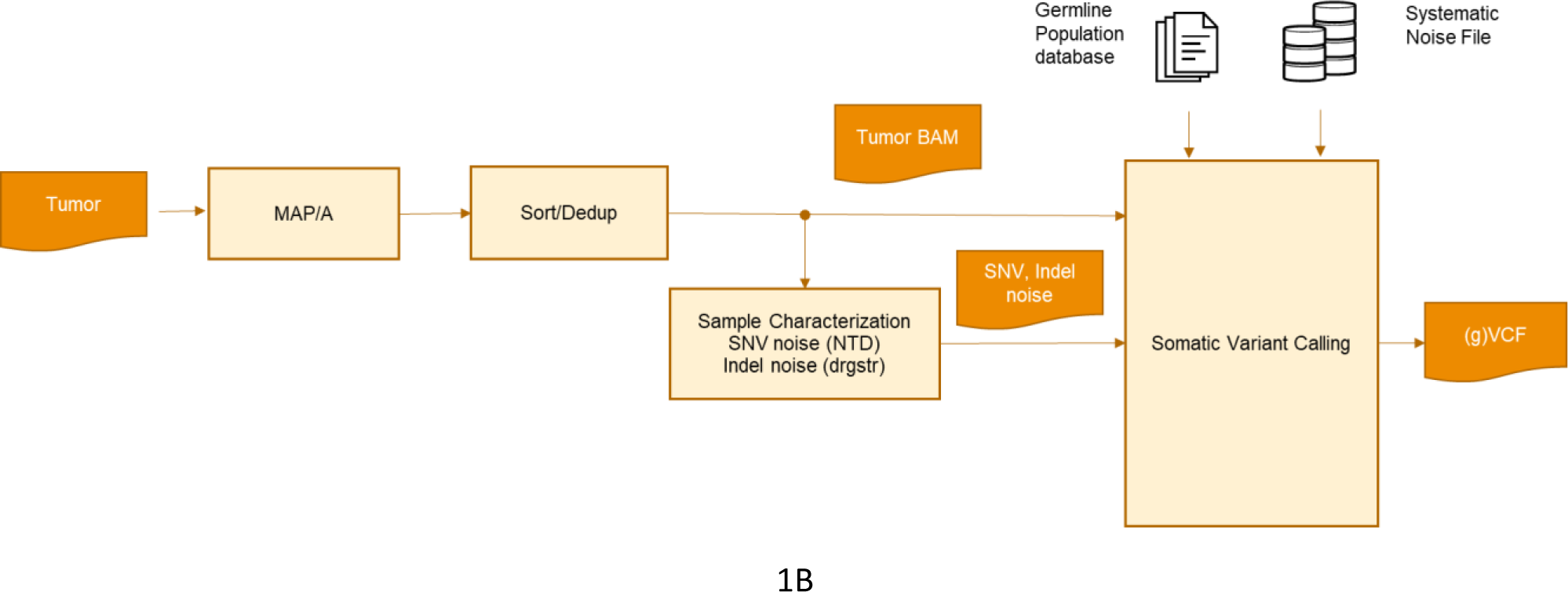
DRAGEN end-to-end somatic pipelines: (A) tumor/normal pipeline; (B) tumor-only pipeline.

An ideal bioinformatics pipeline enables seamless end-to-end processing of any sequencing reads generated from different sample inputs, library prep protocols, and sequencing configurations. To achieve this goal, DRAGEN offers a complete solution to map and align raw sequencing reads to the reference genome, sort the reads, identify polymerase chain reaction (PCR) duplicates, suppress sequencing errors in the presence or absence of unique molecular identifiers (UMIs), detect somatic SNVs and indels, and provide a somatic quality score (SQ) that can be leveraged to filter low confidence variants and reduce false positives (FP)s (Fig. 1).

As shown in Fig. 1, DRAGEN includes a sample-specific characterization step, which takes as input the aligned BAM (Binary Alignment and Map), and outputs estimates of SNV nucleotide error bias and indel error rates, both of which then inform the parameters for the hidden Markov model (HMM) that performs the read likelihood calculation inside the somatic variant caller. The SNV Nucleotide (NTD) Error Bias Estimation corrects for sample-wide nucleotide substitution biases, e.g., increased C->T rate due to oxidation/deamination that is common in Formalin-Fixed Paraffin-Embedded (FFPE) samples. The bias is estimated at a subset of sites in the sample and the estimated parameters are used to adjust basecall quality in the HMM. The sample-specific indel error model was first introduced and described in the DRAGEN germline pipeline [8]. It estimates the rates of insertion and deletion errors due to PCR amplification by finding parameters that maximize the probability of producing a set of observed read pileups. Those parameters depend on both the reference short tandem repeat (STR) period and the repeat length. The model optimizes three sets of parameters: 1) the gap open penalties (GOP) directly related to indel error rate; 2) the gap continuation penalties (GCP) related to indel length; and 3) a separate parameter directly related to the probability of indel variants of any non-zero length. In the case of the somatic tumor normal workflow, INDEL error rate estimation modules can be run independently on the tumor and normal reads to characterize them separately.

The normal sample, if present, is used to avoid calls at sites with germline variants or systematic sequencing artifacts. In addition, also shown in Fig. 1A, it is possible to provide the somatic variant caller with a germline population database file and a systematic noise file. While these files are critical in a tumor-only workflow to annotate variants as germline based on population databases (gnomAD and 1000 genomes) and filter systematic noise artifacts, respectively, they can also be optionally used in a tumor-normal workflow to remove any remaining germline and/or systematic noise that escaped the matched normal sample.

## 3 DRAGEN Somatic Variant Caller

### 3.1 DRAGEN somatic variant caller workflow

The DRAGEN somatic variant caller workflow is described in Fig. 2 below. It takes in a sorted and aligned tumor BAM as input, and optionally a sorted and aligned normal BAM. The first step consists of looking for sufficient coverage and evidence of variants in the tumor reads to establish active regions. Since DRAGEN is a haplotype-based variant caller, the reads covering an active region are then locally assembled via a DeBrujin graph to generate a set of candidate haplotypes. This step is similar in concept to GATK4/Mutect2 [1]. Once the haplotypes are assembled, they are aligned against the human reference to identify variants. It is possible to augment the events generated by the graph by recruiting events from “column-wise” detection which consists of counting the number of reads supporting a variant at a given position in a read pileup. The HMM then computes a likelihood for each read-haplotype pair, taking into account the SNV and INDEL sample-specific noise estimates computed upstream of the variant caller. The mutation calling step then calculates the posterior probability of a candidate somatic variant being present at the site, integrating over all possible VAF, and computes SQ It then emits a call in the VCF (or optionally gVCF) if the SQ is above a call threshold. Downstream of mutation calling, a set of filters are applied to remove FPcalls and germline variant tagging can be applied.

**Figure 2.**
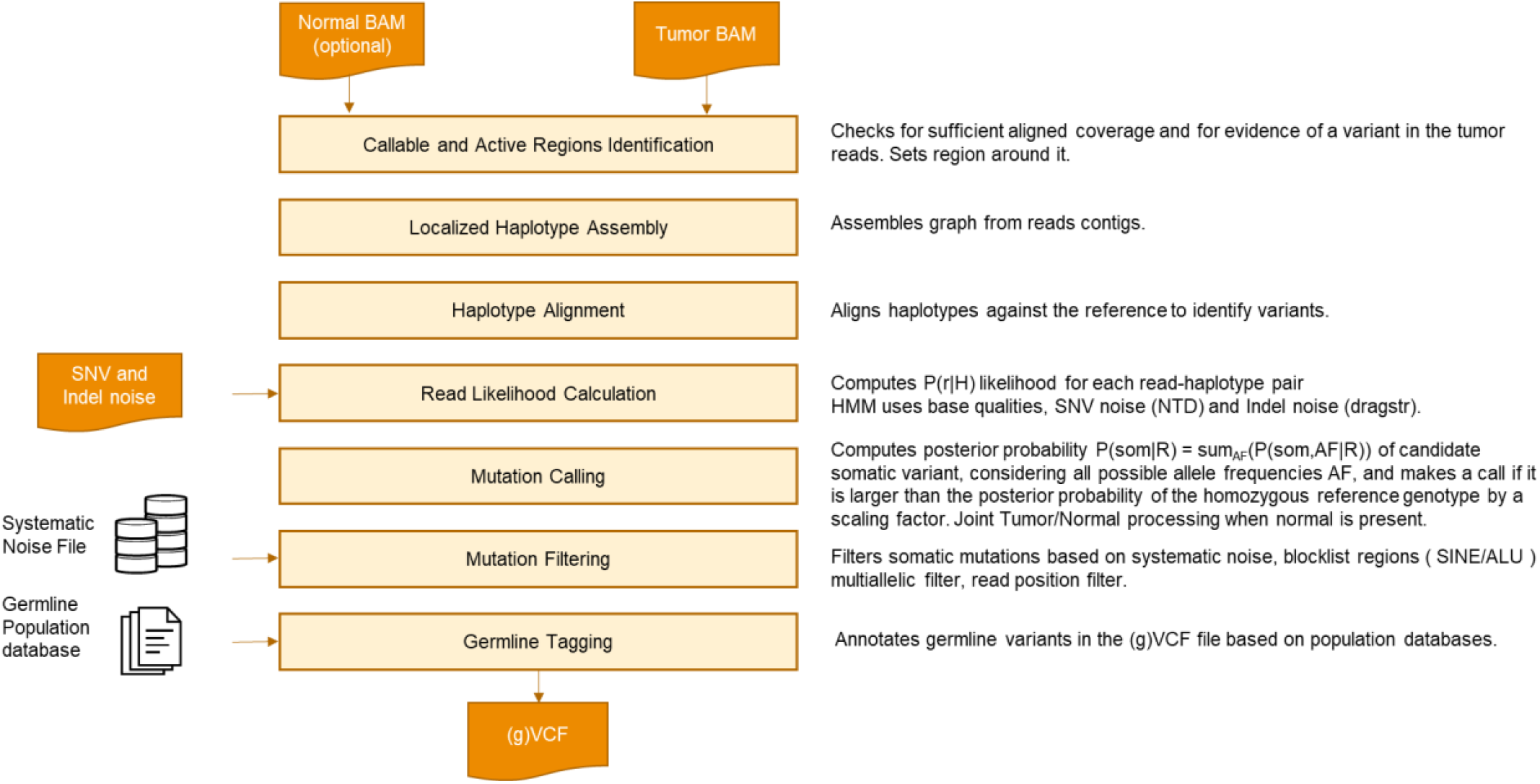
DRAGEN somatic small variant caller

### 3.2 Further details on DRAGEN mutation calling and filtering steps

As outlined in Fig. 3, one of the key differentiators of the DRAGEN mutation calling step is its ability to perform a joint analysis of the tumor and normal samples, in the case of a T/N workflow. While it shares a similar concept with the joint genotyping of Strelka2 [2], the DRAGEN joint T/N analysis incorporates its own probability models of common error modes such as strand bias, orientation bias, and mis-mapping based on existing models originally implemented in the DRAGEN germline pipeline [8].

**Figure 3.**
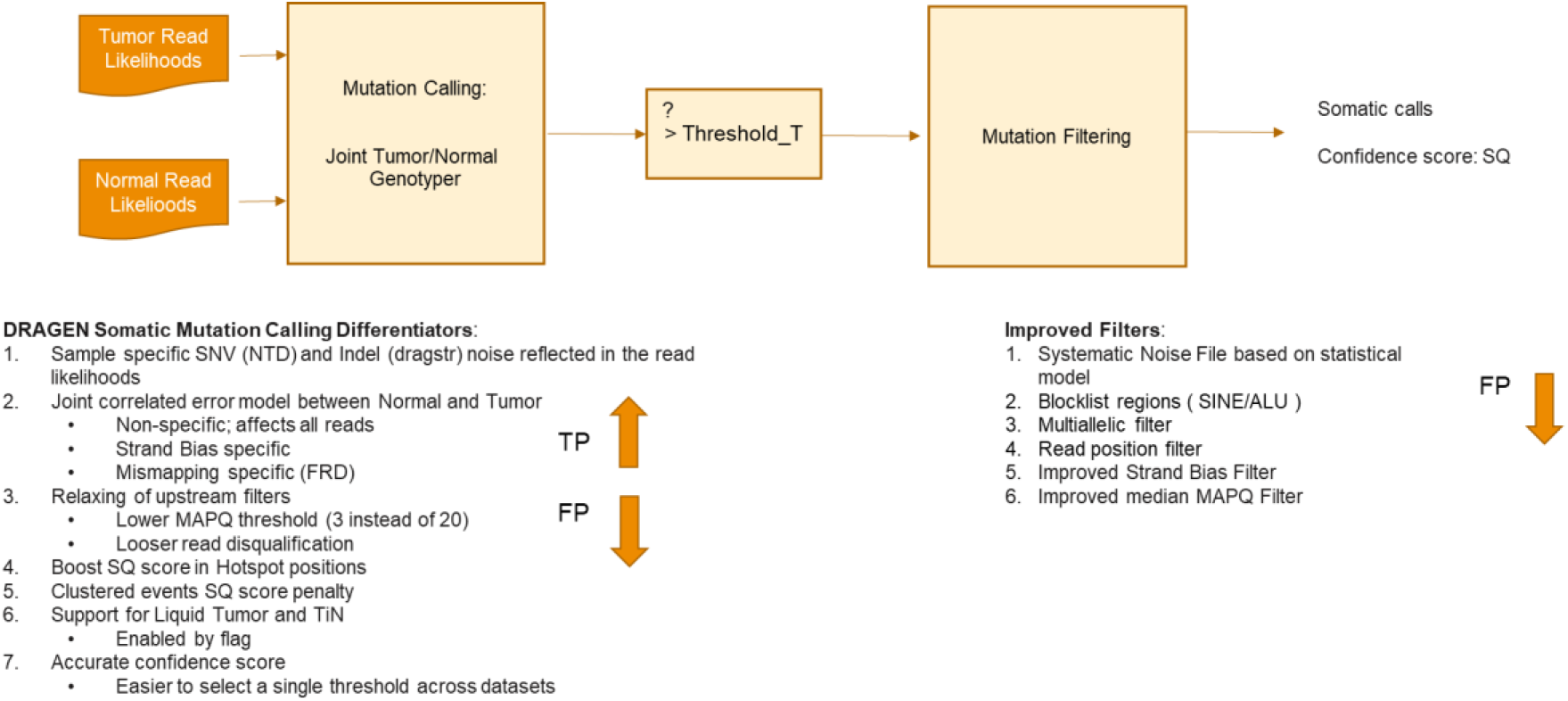
DRAGEN mutation calling and filtering steps

Because of its increased robustness against shared noise between the tumor and normal samples, the DRAGEN variant caller can apply more relaxed thresholds when accepting reads for downstream processing. For example, it takes in reads with MAPQ as low as 3, while other tools such as GATK4/Mutect2, which don’t have such robustness against noise, apply a more stringent MAPQ threshold of 20 to filter out mid-to-low confidence mapped reads (that would otherwise cause FP). An overly high MAPQ threshold can cause valuable evidence of mutations to be lost, hence being able to lower the MAPQ threshold yields increased sensitivity. A similar principle is applied to the read disqualification threshold, where the DRAGEN mutation calling step accepts reads with more mismatches in their alignment than third party tools and leverages the evidence of those reads to increase sensitivity.

Downstream of the mutation calling step, a set of filters may be applied to reduce FP calls. Some filters are based on the position of the mutation within the supporting reads, strand bias, or are related to certain regions of the genome known to be noisy.

### 3.3 Joint analysis of tumor and normal samples

One particularly challenging problem in calling small somatic variants in tumors is to distinguish them from both germline variants and systematic sequencing errors. Although many tools are available to perform somatic variant calling [1–2, 9–14], there is substantial room for improvement in both accuracy and speed. To address this challenge, it is common to sequence matched tumor and normal samples, where a sample from the tumor is accompanied by a normal (e.g., blood) sample from the same patient (Fig. 1A). The hope is that the normal sample will not contain any tumor mutations, although this assumption is violated in the case of Tumor-in-Normal (TiN) contamination. Regardless of whether TiN is present, the normal sample should still be useful for screening out both germline variants and systematic sequencing errors. In the absence of somatic copy number variation these artifacts are expected to be present in both samples at roughly the same frequency, whereas somatic variants (if present in the normal sample at all) should be present at a higher frequency in the tumor sample.

In its joint tumor/normal analysis, DRAGEN uses a Bayesian approach in which the tumor and normal allele frequencies are treated as dependent random variables. The core of the model is a pair of joint prior distributions of the allele frequencies for the candidate somatic allele in the tumor and normal samples: one joint distribution for the somatic variant hypothesis (Fig. 4A) and one for the corresponding non-somatic hypothesis (Fig. 4B). This part of the model is similar in concept to the shared error model in Strelka2 [2]. The caller will favor the somatic or non-somatic hypothesis depending on which of these two joint allele frequency distributions represents a better fit to the data in the two samples. By analyzing both samples jointly, the model can account for systematic error patterns that are shared between the samples and for TiN contamination, as is common in liquid tumors. To account for TiN contamination, DRAGEN uses a maximum contamination tolerance parameter and, in contrast to alternative approaches such as DeTiN [15], does not require an explicit estimate of the contamination level.

**Figure 4.**
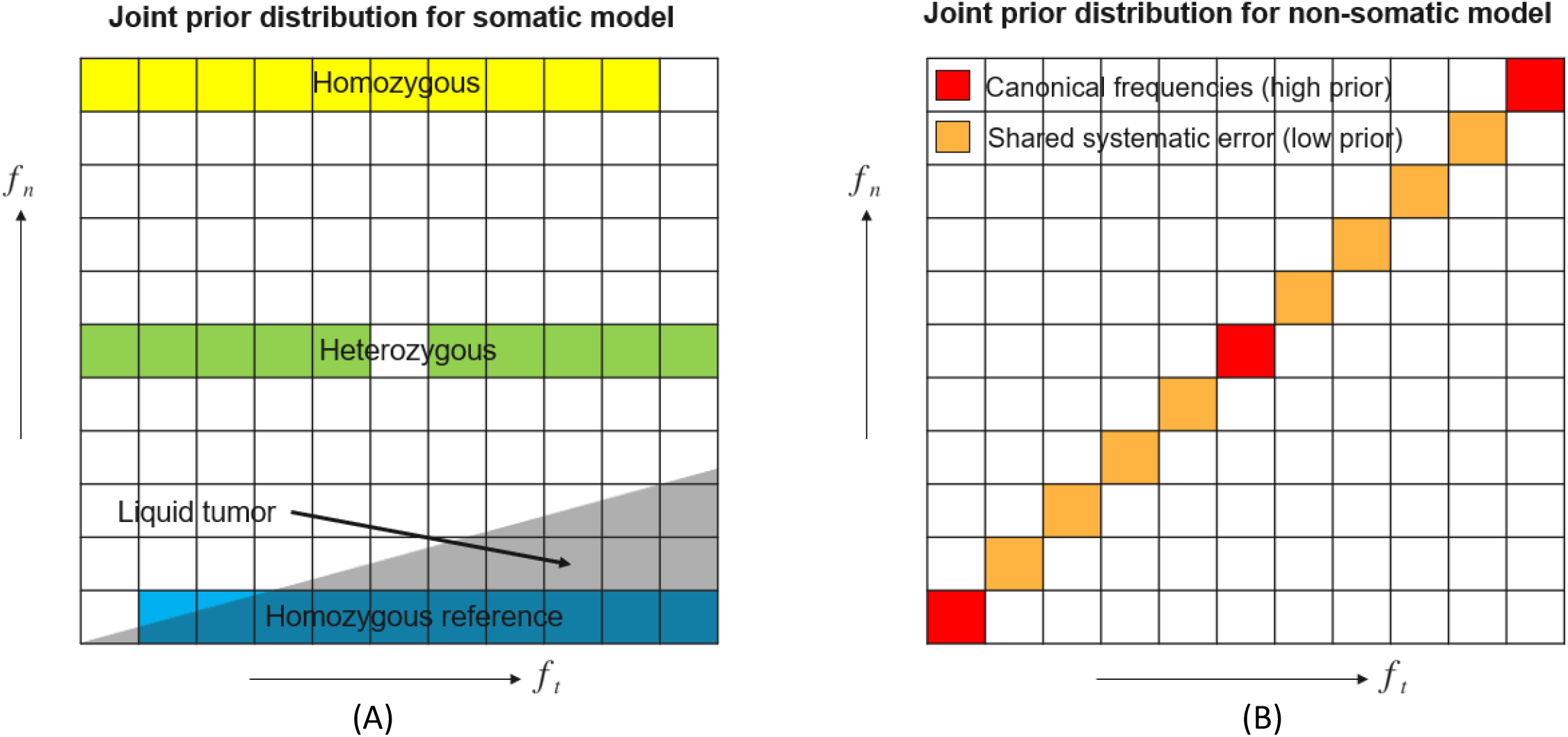
Joint distributions of the frequency of a candidate somatic allele in the tumor and normal samples for each of the two models. (A) Somatic model: the normal allele frequency (fn) is expected to be one of the canonical frequencies 0, 0.5, or 1, while the tumor allele frequency (ft) is any value different from the normal frequency. In the liquid tumor model the triangular component (with the diagonal line having a slope) is added, allowing the alt allele frequency in the normal sample to increase up to a fraction of the alt frequency in the tumor sample despite the germline genotype being homozygous reference. (B) Non-somatic model: the tumor and normal frequencies are equal to each other and is normally at one of the canonical frequencies. However, there is a small probability that systematic error will be present, yielding a non-canonical allele frequency in both samples.

The shared error model accounts for systematic error patterns, such as sequencing, mapping, and alignment errors, potentially affecting all reads in both samples. In addition, it allows strand-specific shared error (potentially affecting the reads on only one strand in both samples) and mis-mapping shared error (potentially affecting only the reads below a variable mapping quality threshold in both samples). The model for mis-mapping error is inspired by the Foreign Read Detection (FRD) model in the DRAGEN germline pipeline [6] and treats the distance to the second-closest reference sequence (from which mis-mapped reads are most likely to originate) as a random variable that governs the mapping quality threshold.

### 3.4 Hypotheses and posterior probability calculation

At every locus where candidate variant alleles have been proposed, the DRAGEN genotyper considers each candidate variant allele in isolation. The main task of the genotyper is to decide whether the candidate somatic variant is (a) a real somatic variant (described by the somatic model); (b) a real germline variant (described by the clean case of the non-somatic model); or (c) just noise (either systematic noise, described by one of the noisy cases of the non-somatic model, or independent per-read noise that can be present in any of the models). These possible explanations of the observed reads correspond to different hypotheses for which probabilities are to be calculated.

We refer to the candidate variant allele as the “somatic” allele, even when discussing hypotheses in which it appears as a germline variant or a noise allele rather than a somatic variant. When an indel variant is present, it is common for independent sequencing errors to result in reads supporting alleles of different lengths, leading to multi-allelic loci. While considering a candidate variant allele, any other allele with enough supporting reads in the normal sample is considered a “non-somatic” allele. Besides the reference allele, non-somatic alleles could be real germline variants, noise alleles, or real somatic alleles that are not the candidate somatic allele currently under consideration. In cases where there are multiple non-somatic alleles, all of the non-somatic alleles are considered together as if they were a single non-somatic allele, so that we only ever consider hypotheses involving two alleles. The resulting two-allele model leads to a simplified calculation at multi-allelic loci.

Given a somatic allele, DRAGEN calculates posterior probabilities for a range of genotype hypotheses, under the assumption that the normal sample conforms to the diploid germline genotype while the tumor sample is a mixture of the germline genotype and, if present, the somatic allele. The two-allele model yields three hypotheses for the germline genotype: *G*_*g*_ ∈ {*g*_*ref*_, *g*_*het*_, *g*_*hom*_}, referring to homozygous non-somatic, heterozygous, and homozygous, respectively. For the somatic genotype web consider only two states: *G*_*s*_ ∈ {*g*_*nonsom*_, *g*_*som*_}, referring to the absence and presence of the somatic variant in the tumor sample. When the somatic variant is absent, *G*_*s*_ = *g*_*nonsom*_, we further distinguish between sub-cases representing different error modes. We treat the state where a somatic variant is present, *G*_*s*_ = *g*_*som*_, as free of systematic error by definition to avoid false positive somatic calls when there is evidence of systematic error.

For each joint hypothesis on the tumor and normal samples, we calculate the posterior probability:

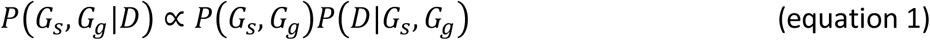

where *D* refers to the full set of reads from both samples. For the likelihood term, *P*(*D*|*G*_*s*_, *G*_*g*_), we use either the non-somatic model (*G*_*s*_ = *g*_*nonsom*_) or the somatic model (*G*_*s*_ = *g*_*som*_). Within each of these models, different subsets of reads (distinguished by sample, strand, and/or mapping quality) are assigned different expected allele frequency distributions. The likelihood for each read subset is then expanded as an integral over allele frequencies. The prior probability term *P*(*G*_*s*_, *G*_*g*_) assumes independence between the germline and somatic priors. The final variant score is the phred-scale posterior probability calculated in equation 1.

## 4 Somatic WGS T/N Results

### 4.1 Benchmarking datasets

We benchmarked DRAGEN on publicly available datasets from four cancer cell lines with curated truth sets [16, 19]. We chose the NYGC and SEQCII datasets because they were created and curated by third parties, to ensure an impartial assessment. The somatic variants in the NYGC cancer cell lines were characterized by sequencing them deeper than usual, on HiSeqX and Novaseq [16]. Several callers were run and “High confidence” variants were defined by the consensus calls (2+ callers for SNV). The high-confidence somatic SNVs and indels of the HCC1395 SEQCII cell line were obtained based primarily on 21 pairs of tumor-normal WGS replicates from six sequencing centers; sequencing depth ranged from 50X to 100X [20, 28]. The reference call set of somatic mutations was generated based on consensus across sequencing sites and mutation calling pipelines [7, 20]. Because multiple replicates of each of the HCC1395 (tumor) and HCC1395BL (normal) cell lines are available, it is possible to mix in-silico tumor and normal reads to achieve various tumor purity levels. As such we generated two tumor-normal pairs with 75% and 20% tumor purity levels to simulate high and low tumor purity samples.

For the cancer cell lines, we used the public truth sets to measure sensitivity and false negative counts, but we do not trust the truth sets to be sufficiently complete to produce reliable false positive counts. We therefore measure false positive counts and precision on matching normal-normal datasets that have been selected to match the tumor-normal datasets in terms of sequencing platform, sample preparation chemistry, and average coverage. For the precision/FP count measurements of the SEQCII datasets, we generated one normal-normal dataset from the HCC1395 normal cell lines. For the precision/FP count measurements of the NYGC datasets, we generated one normal-normal dataset from the HCC1143 normal cell line (and used it for both HCC1143 and HCC1187 T/N pairs), and one normal-normal dataset from the COLO829 normal cell line, used for the COLO829 T/N pair.

Sensitivity on the cancer cell lines depends strongly on the allele frequencies of the variants in the truth set, as the datasets do not have sufficient coverage (80x on the tumor) to confidently detect variants with allele frequency below 5%. We therefore give details of the VAF ranges of the truth sets in Table 2.

In addition to cancer cell lines, we internally generated germline admixture datasets with a range of tumor purity and tumor-in-normal contamination levels (Table 1). The admixture datasets were created from germline BAMs of NA12877 and NA12878 [22] so we could explore the effects of tumor purity and tumor-in-normal contamination under controlled conditions.

**Table 1.** Datasets for benchmarking. Cancer cell lines datasets are publicly available with independently curated truth sets. Admixture datasets were created by spiking reads from a germline sample of NA12878 into a dataset from NA12877, with different levels of tumor purity and tumor-in-normal (TiN) contamination.

| Dataset | Type | Depth (T:N) | Total true variants |
| --- | --- | --- | --- |
| HCC1395 SRR7890895 75pc | Breast carcinoma, 75% dilution | 82:45 | 41482 |
| HCC1395 SRR7890902 20pc | Breast carcinoma, 20% dilution | 89:45 | 41482 |
| Normal/Normal (HCC1395) | Matched Normal | 85/45 | 0 |
| NYGC COLO829 | Melanoma | 81:42 | 38386 |
| Normal/Normal (COLO829) | Matched Normal | 72/36 | 0 |
| NYGC HCC1143 | Breast carcinoma | 83:42 | 25183 |
| NYGC HCC1187 | Breast carcinoma | 79:40 | 13542 |
| Normal/Normal (HCC1143) | Matched Normal | 73/37 | 0 |
| AdmixNovaPur80Tin0 | Synthetic, Tumor purity 80%, TiN contamination 0% | 109:40 | 1460397 |
| AdmixNovaPur50Tin0 | Synthetic, Tumor purity 50%, TiN contamination 0% | 110:40 | 1460397 |
| AdmixNovaPur20Tin0 | Synthetic, Tumor purity 20%, TiN contamination 0% | 107:40 | 1460397 |
| AdmixNovaPur80Tin5 | Synthetic, Tumor purity 80%, TiN contamination 5% | 108:41 | 1460397 |
| AdmixNovaPur80Tin20 | Synthetic, Tumor purity 80%, TiN contamination 20% | 108:42 | 1460397 |
| AdmixNovaPur40Tin5 | Synthetic, Tumor purity 40%, TiN contamination 5% | 109:41 | 1460397 |
| AdmixNovaPur40Tin20 | Synthetic, Tumor purity 40%, TiN contamination 20% | 109:42 | 1460397 |

### 4.2 Somatic variant calling accuracy

In this section, we show the results on the somatic WGS T/N small variant calling accuracy across the three callers DRAGEN, Mutect2, and Strelka2.

Even though it is not shown here, the accuracy of the DRAGEN™ 4.0 tumor-only somatic small-variant caller was also recently validated in the “NCTR Indel Calling from Oncopanel Sequencing Data Challenge” [25, 26], where it won top place in precision and F1 score in the applicability challenge (Panel X). DRAGEN also showed high and consistent accuracy across the three oncopanels (A, B, and X) by earning the best F1 score when averaged across all three panels.

For the results presented here, we collected the somatic WGS T/N small variant calling accuracy on three metrics: total number of errors (FP+FN), sensitivity, and precision, for both SNV and indel separately.

Fig. 5 compares the SNV and indel calling accuracy of DRAGEN, Strelka2, and Mutect2 on the NYGC cell lines, COLO829, HCC1143, and HCC1187.

**Figure 5.**
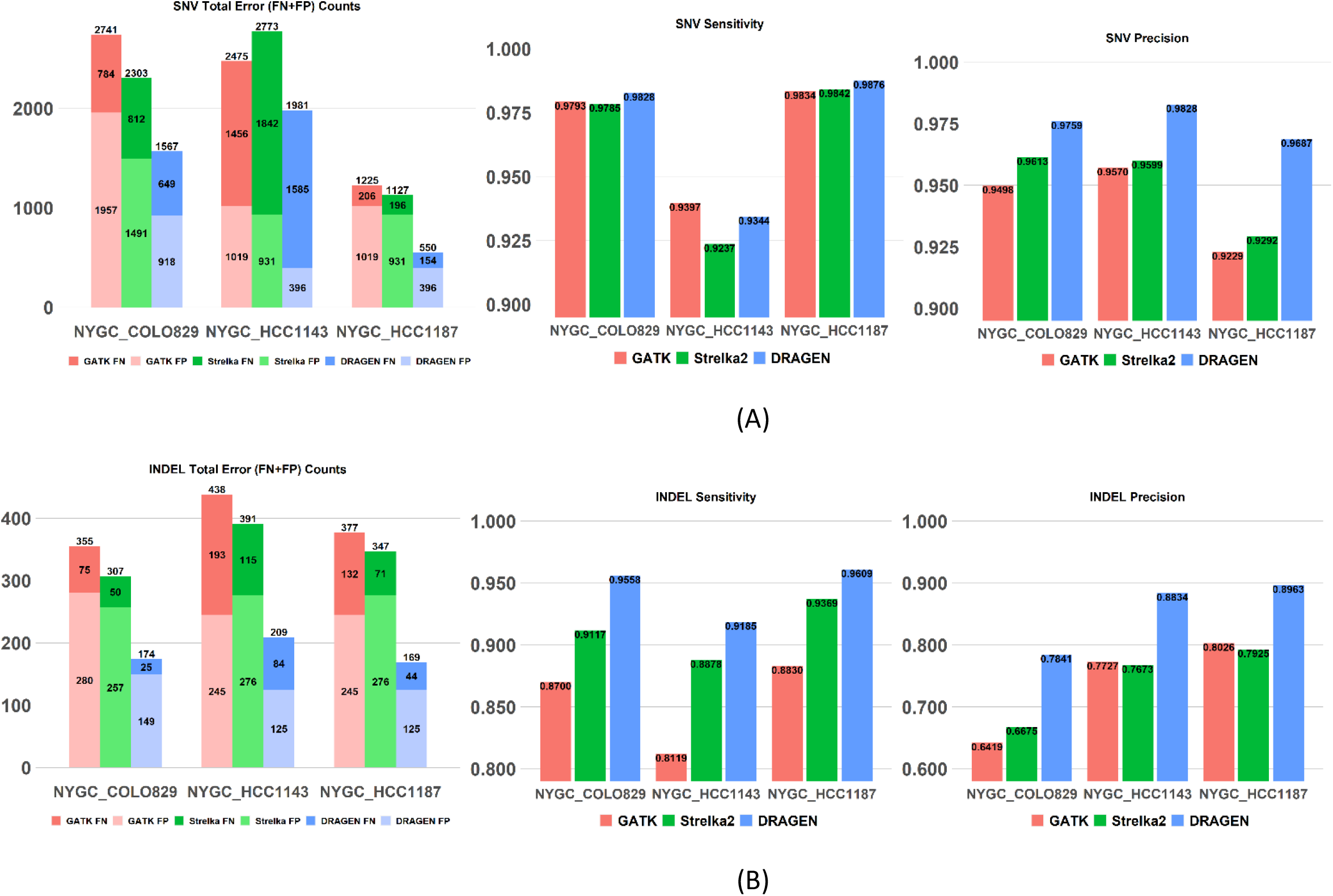
SNV and indel accuracy comparison on the NYGC cell lines (COLO829, HCC1143, HCC1187), between DRAGEN, Strelka2, and Mutect2. A) SNV results B) indel results.

For both SNV and indel, DRAGEN yields a substantial reduction in the total number of errors (FP+FN) compared to the other two tools, with a substantial drop in the number of FP calls. DRAGEN brings ~40-60% SNV and indel FP count reduction compared to the other two tools, yielding 0.13-0.31 SNV FP/MB. The FP rate is higher for the COLO829 cell line compared to HCC1143/1187, but that is a consistent finding across all three tools, so it is most likely sample specific. Please note the FP count is the same for all three tools between HCC1143 and HCC1187 because we used the same normal-normal pair for the FP count and precision measurements.

In terms of sensitivity, DRAGEN brings ~20% SNV FN reduction for COLO829 and HCC1187, the gains are smaller for HCC1143, likely because of the lower VAF distribution for this cell line (cf. Table 2). The gains in indel FN are striking, ranging between 30-67% indel FN reduction compared to the other two tools.

**Table 2.** Factors limiting sensitivity in cancer cell line datasets. Sensitivity decreases with the number of variants abovethe 5%VAF, or as tumor purity drops (causing a general decrease in VAF), or astumor-in-normal contamination increases.

| Dataset | Truth set VAF range | % truth set variants above 5% VAF | SNV sensitivity | SNV FN | Indel sensitivity | Indel FN |
| --- | --- | --- | --- | --- | --- | --- |
| HCC1395 SRR7890895 75pc | 0.015 – 0.75 | 92.64 | 0.93641 | 2508 | 0.906135 | 153 |
| HCC1395 SRR7890902 20pc | 0.004 – 0.2 | 61.78 | 0.700489 | 11813 | 0.633742 | 597 |
| NYGC COLO829 | 0.0151 – 1.0 | 98.58 | 0.982837 | 649 | 0.95583 | 25 |
| NYGC HCC1143 | 0.0059 – 1.0 | 94.51 | 0.934358 | 1585 | 0.918526 | 84 |
| NYGC HCC1187 | 0.0253 – 1.0 | 99.05 | 0.987593 | 154 | 0.960854 | 44 |

**Table 3.** Comparison of combined SNV and indel FP count between DRAGEN, Mutect2, and Strelka2 on the NYGC cell lines, and DRAGEN gains in FP reduction vs. the other two tools

|  | Dataset | DRAGEN<br>FP count | Strelka2<br>FP count | Mutect2<br>FP count | DRAGEN<br>FP/MB | Strelka2<br>FP/MB | Mutect2<br>FP/MB | DRAGEN gain<br>vs Strelka2 | DRAGEN gain<br>vs Mutect2 |
| --- | --- | --- | --- | --- | --- | --- | --- | --- | --- |
| SNV | NYGC COLO829 | 918 | 1491 | 1957 | 0.31 | 0.50 | 0.65 | 0.38 | 0.53 |
|  | NYGC<br>HCC1143/HCC1187 | 396 | 931 | 1019 | 0.13 | 0.31 | 0.34 | 0.57 | 0.61 |
| indel | NYGC COLO829 | 149 | 257 | 280 | 0.05 | 0.09 | 0.09 | 0.42 | 0.47 |
|  | NYGC<br>HCC1143/HCC1187 | 125 | 276 | 245 | 0.04 | 0.09 | 0.08 | 0.55 | 0.49 |

**Table 4.** Comparison of SNV and indel FN count between DRAGEN, Mutect2, and Strelka2 on the NYGC cell lines, and DRAGEN gains in FN reduction vs. the other two tools

|  | Dataset | DRAGEN<br>FN count | Strelka2<br>FN count | Mutect2<br>FN count | DRAGEN gain<br>vs Strelka2 | DRAGEN gain<br>vs Mutect2 |
| --- | --- | --- | --- | --- | --- | --- |
| SNV | NYGC COLO829 | 649 | 812 | 784 | 0.20 | 0.17 |
|  | NYGC HCC1143 | 1585 | 1842 | 1456 | 0.14 | -0.09 |
|  | NYGC HCC1187 | 154 | 196 | 206 | 0.21 | 0.25 |
| indel | NYGC COLO829 | 25 | 50 | 75 | 0.50 | 0.67 |
|  | NYGC HCC1143 | 84 | 115 | 193 | 0.27 | 0.56 |
|  | NYGC HCC1187 | 44 | 71 | 132 | 0.38 | 0.67 |

**Table 5.** Comparison of SNV and indel FP count between DRAGEN, Mutect2, and Strelka2, and DRAGEN gains in FP reduction vs. the other two tools

|  | Dataset | DRAGEN<br>FP count | Strelka2<br>FP count | Mutect2<br>FP count | DRAGEN<br>FP/MB | Strelka2<br>FP/MB | Mutect2<br>FP/MB | DRAGEN gain<br>vs Strelka2 | DRAGEN gain<br>vs Mutect2 |
| --- | --- | --- | --- | --- | --- | --- | --- | --- | --- |
| SNV | HCC1395 75%/20% | 465 | 905 | 3703 | 0.16 | 0.30 | 1.23 | 0.49 | 0.87 |
| indel | HCC1395 75%/20% | 204 | 198 | 1339 | 0.07 | 0.07 | 0.45 | -0.03 | 0.85 |

**Table 6.** Comparison of SNV and indel FN count between DRAGEN, Mutect2, and Strelka2 on the SEQCII cell lines, and DRAGEN gains in FN reduction vs. the other two tools

|  | Dataset | DRAGEN<br>FN count | Strelka2<br>FN count | Mutect2<br>FN count | DRAGEN gain<br>vs Strelka2 | DRAGEN gain<br>vs Mutect2 |
| --- | --- | --- | --- | --- | --- | --- |
| SNV | HCC1395 75% | 2508 | 2580 | 3408 | 0.03 | 0.26 |
|  | HCC1395 20% | 11813 | 12231 | 12057 | 0.03 | 0.02 |
| indel | HCC1395 75% | 153 | 207 | 285 | 0.26 | 0.46 |
|  | HCC1395 20% | 597 | 865 | 675 | 0.31 | 0.12 |

Fig. 6 compares the SNV and indel calling accuracy of DRAGEN, Strelka2, and Mutect2 on the SEQCII cell lines, at two different tumor purity levels: HCC1395_75% and HCC1395_20%.

**Figure 6.**
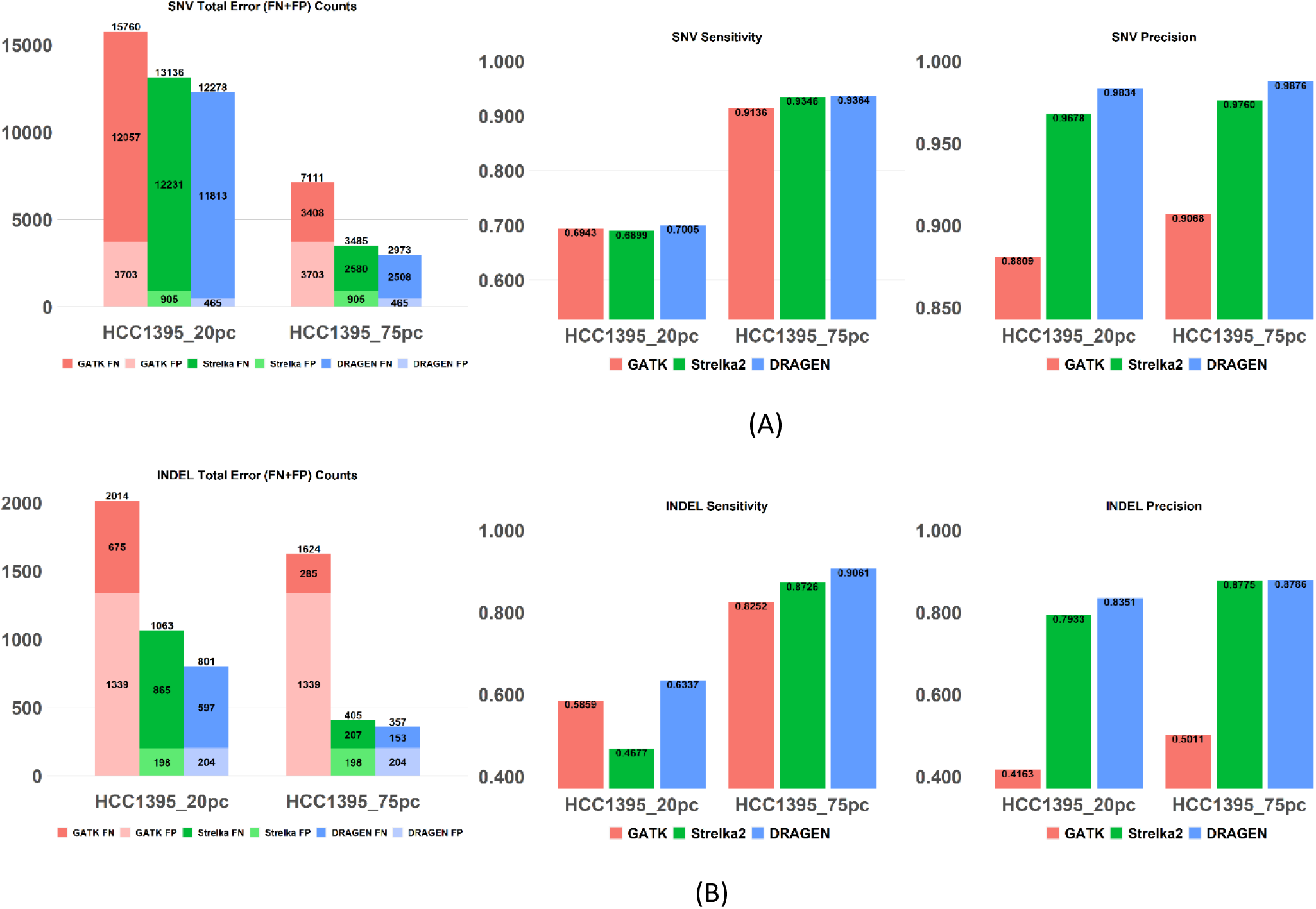
SNV and indel accuracy comparison on the SEQCII cell lines at two different tumor purity levels (HCC1395_75% and HCC1395_20%), between DRAGEN, Strelka2, and Mutect2. A) SNV results B) indel results.

For the SEQCII cell line, both DRAGEN and Strelka2 provide large FP reduction gain vs. Mutect2, suggesting the presence of shared noise error between the normal and the tumor sample for this cell line. Both DRAGEN and Strelka2 process the tumor and normal sample jointly but Mutect2 does not. DRAGEN provides 85% FP reduction compared to Mutect2. The FP rate in indel is comparable between DRAGEN and Strelka2, but DRAGEN achieves substantial SNV FP gains (50% reduction) over Strelka2.

As expected, the SNV sensitivity is much lower in the lower purity tumor (20%), and it is comparable across all three callers (~70% recall). For the high purity tumor, DRAGEN and Strelka2 are comparable and provide a sizeable gain of 26% FN reduction vs. Mutect2.

DRAGEN remains superior in indel recall, for both low and high tumor purity with FN reduction gains ranging from 12-46% compared to the other two tools. These results show the robustness of the DRAGEN somatic variant caller even at lower purity levels: e.g., at 20% tumor purity level, DRAGEN yields 70% and 63% SNV and INDEL sensitivity, respectively, while still maintaining precision at 98.34 and 83.5%, respectively.

Fig. 7 compares the accuracy across the three somatic variant callers on the germline admixtures without tumor-in-normal contamination at different levels of tumor purity. The high false negative count obtained by GATK4/Mutect2 on germline admixture data is largely due to the clustered event filter in this pipeline, which is overly aggressive when the somatic variant rate is high (as is expected in germline admixtures where the simulated somatic variant rate is roughly equal to the germline variant rate). For this experiment, the big picture is easier to understand by focusing on the sensitivity and precision measurements, tabulated in Table 7.

**Figure 7.**
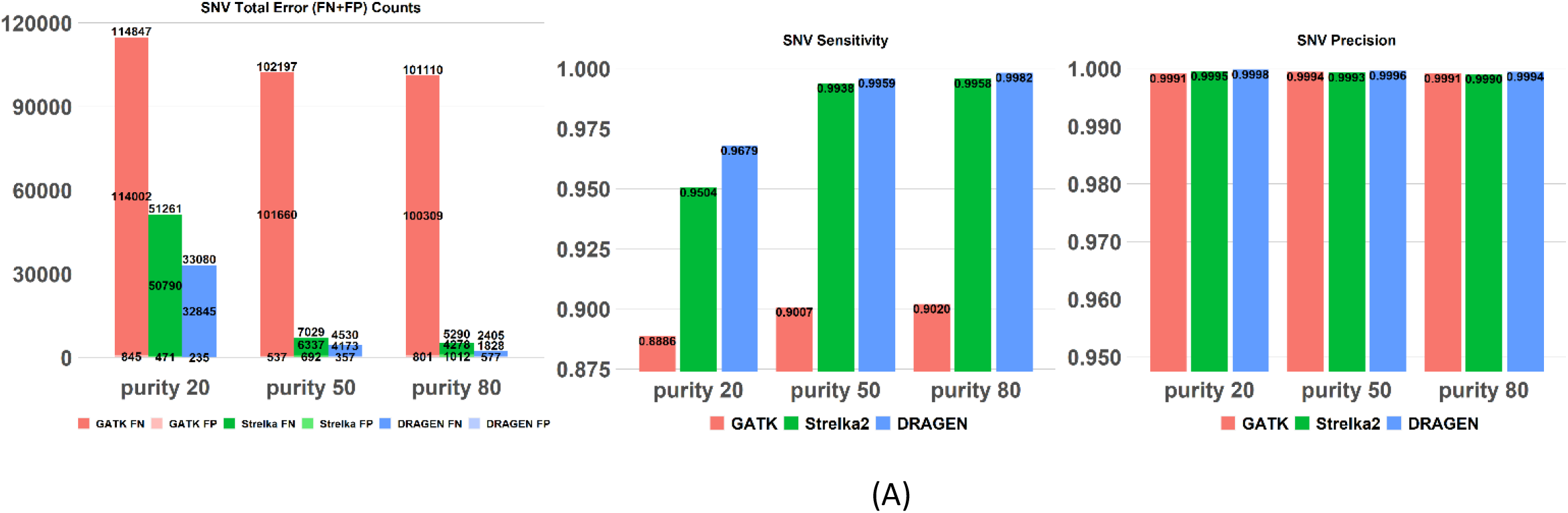
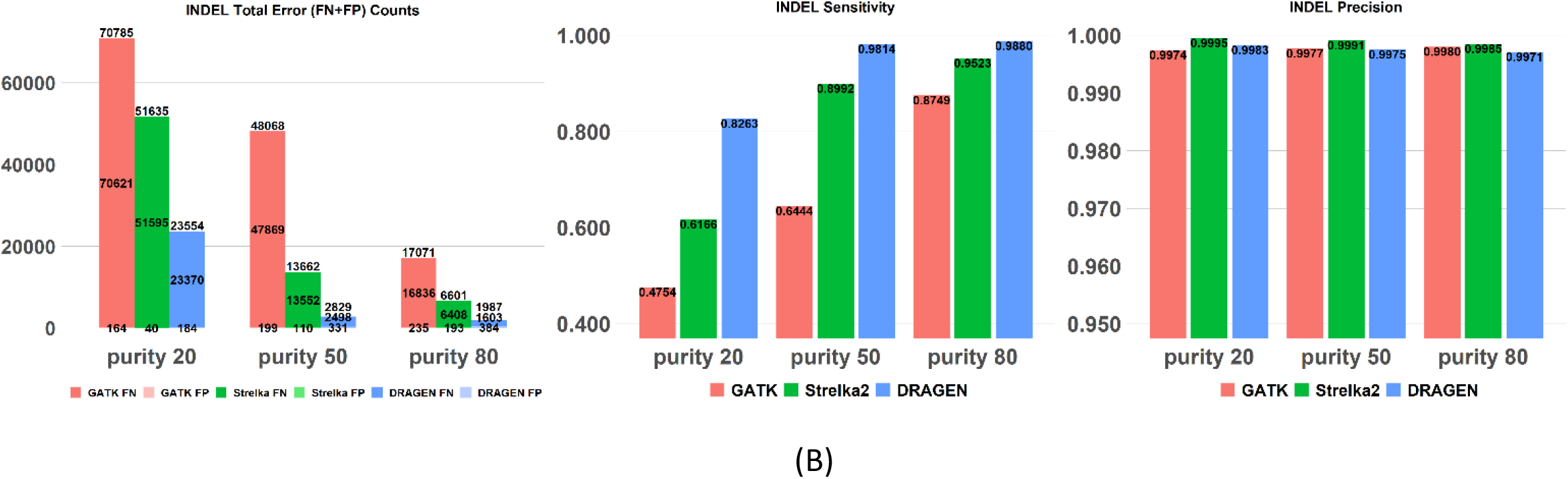
Admixture datasets without TiN contamination, comparing DRAGEN, Strelka2, and Mutect2 at tumor purity levels of 20%, 50%, and 80%.

**Table 7.**
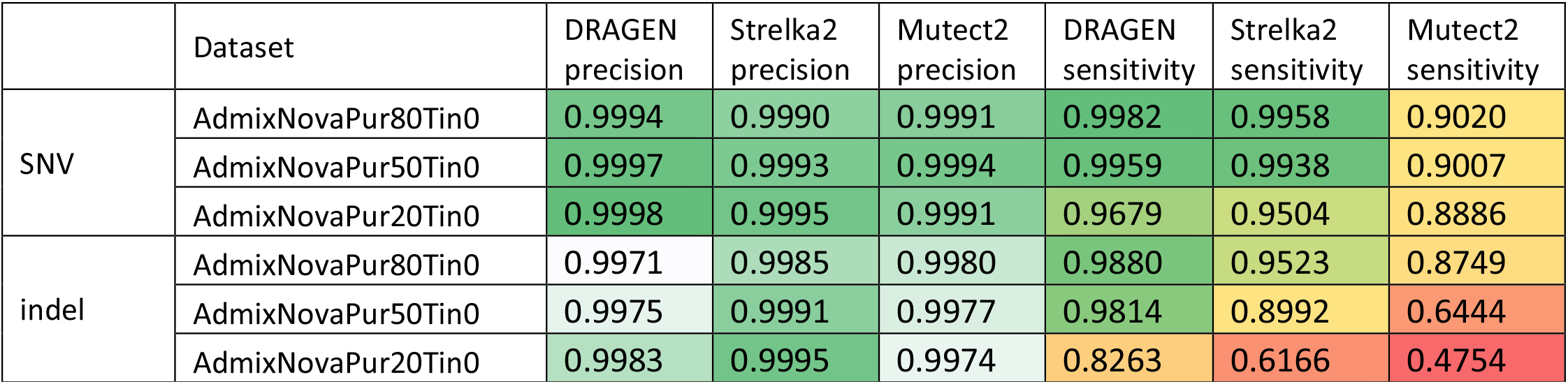
Admixture datasets without TiN contamination, comparing sensitivity and precision of DRAGEN, Strelka2, and Mutect2 at tumor purity levels of 20%, 50%, and 80%.

|  | Dataset | DRAGEN precision | Strelka2 precision | Mutect2 precision | DRAGEN sensitivity | Strelka2 sensitivity | Mutect2 sensitivity |
| --- | --- | --- | --- | --- | --- | --- | --- |
| SNV | AdmixNovaPur80Tin0 | 0.9994 | 0.9990 | 0.9991 | 0.9982 | 0.9958 | 0.9020 |
|  | AdmixNovaPur50Tin0 | 0.9997 | 0.9993 | 0.9994 | 0.9959 | 0.9938 | 0.9007 |
|  | AdmixNovaPur20Tin0 | 0.9998 | 0.9995 | 0.9991 | 0.9679 | 0.9504 | 0.8886 |
| indel | AdmixNovaPur80Tin0 | 0.9971 | 0.9985 | 0.9980 | 0.9880 | 0.9523 | 0.8749 |
|  | AdmixNovaPur50Tin0 | 0.9975 | 0.9991 | 0.9977 | 0.9814 | 0.8992 | 0.6444 |
|  | AdmixNovaPur20Tin0 | 0.9983 | 0.9995 | 0.9974 | 0.8263 | 0.6166 | 0.4754 |

**Table 8.**
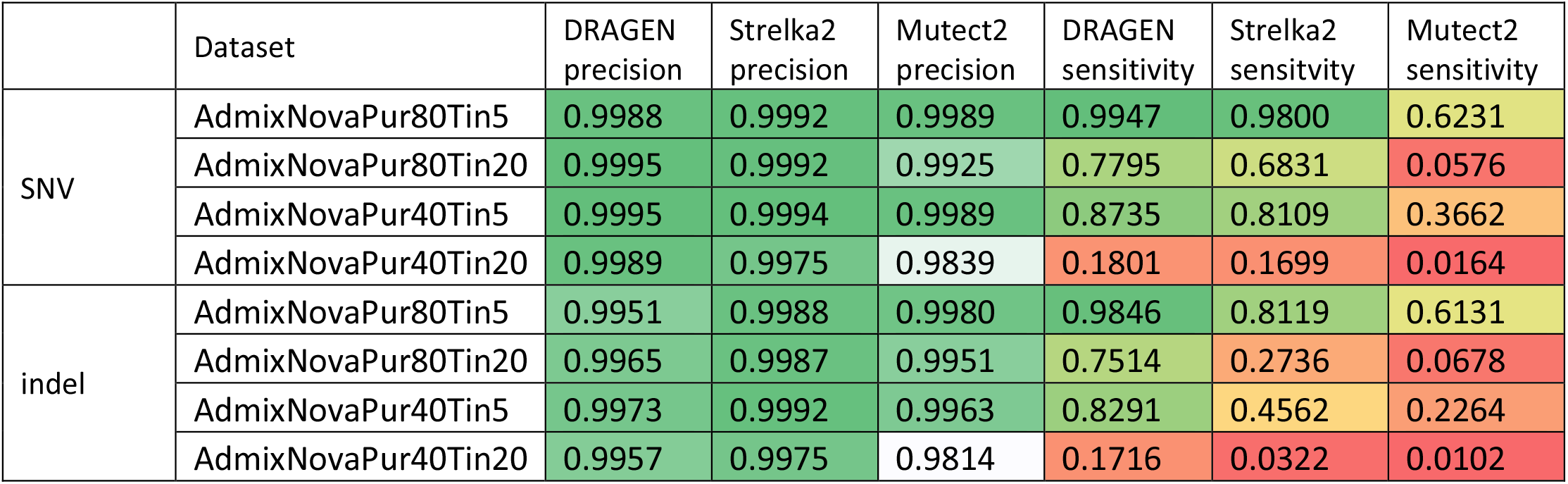
Admixture datasets with TiN contamination of 5% and 20%, comparing sensitivity and precision of DRAGEN, Strelka2, and Mutect2 at tumor purity levels of 40% and 80%.

Table 7 illustrates that all three callers yield high SNV precision regardless of tumor purity with the precision being slightly lower for indels. For sensitivity, DRAGEN yields the best results, especially as the tumor purity decreases, demonstrating the robustness of the DRAGEN somatic variant caller even at lower purity level: e.g., at 20% tumor purity level, DRAGEN yields 96.79% and 82.63% SNV and INDEL sensitivity, respectively, while still maintaining precision ~= 99.9%. Similar to what was found with the cell line datasets, DRAGEN has superior indel performance over the other two callers.

Finally, Fig. 8 and 9 show the impact of TiN contamination of 5% and 20%, respectively, in germline admixtures at different levels of tumor purity. This type of contamination tends to suppress calls if unaccounted for, resulting in large numbers of false negatives for Mutect2, which was not designed to tolerate this type of contamination. Strelka2 does take TiN contamination into account, but DRAGEN nevertheless obtains far better sensitivity, especially for indels. The case with 40% purity and 20% TiN contamination was included as a near-impossible problem to explore the limits of the algorithm – here the allele frequencies in the normal sample are 50% of those in the tumor sample, which is not a condition under which one would expect to be able to detect many true variants. This leads to all of the methods showing a large sensitivity degradation, but DRAGEN nevertheless outperforms the other methods.

**Figure 8.**
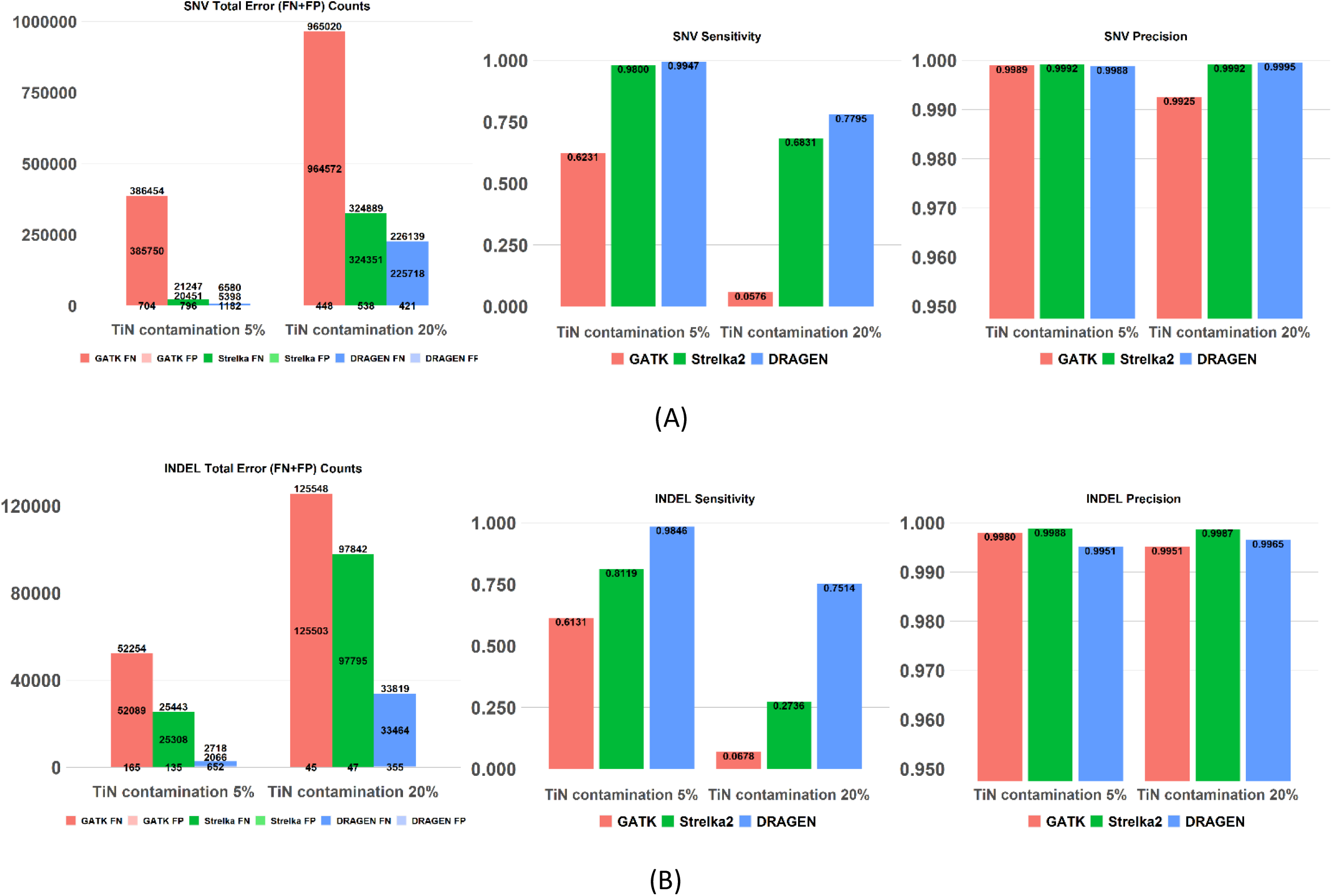
Admixture datasets with 80% tumor purity and 5% or 20% TiN contamination, comparing DRAGEN (liquid tumor mode), Mutect2 and Strelka2.

**Figure 9.**
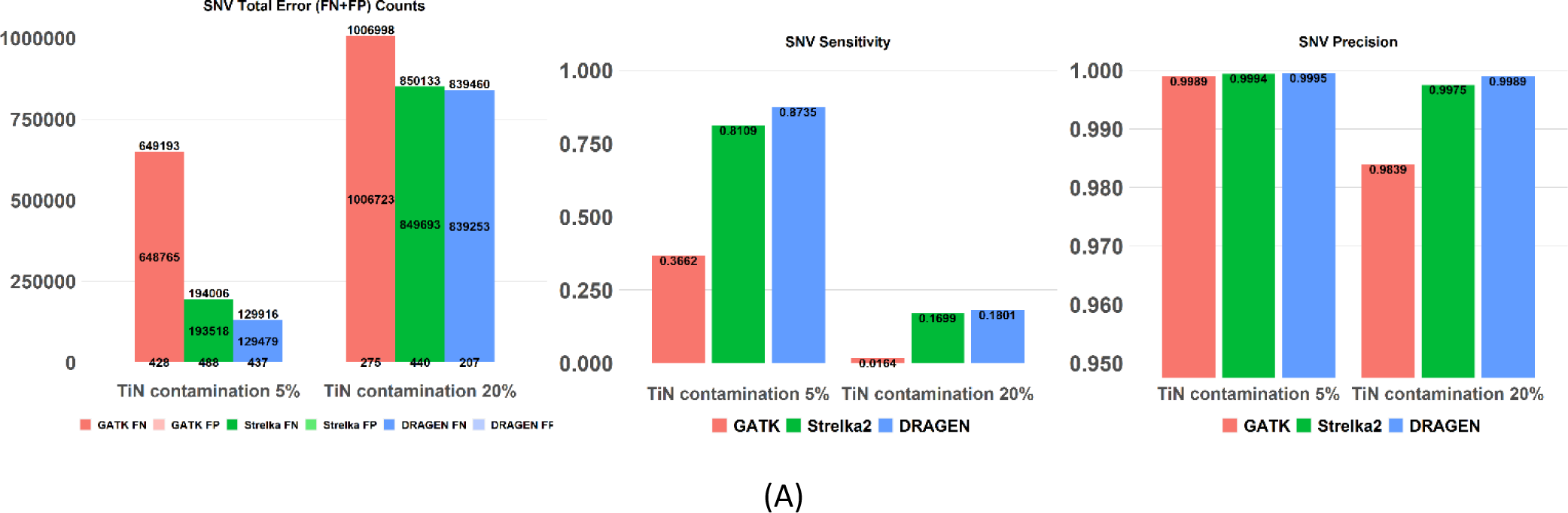
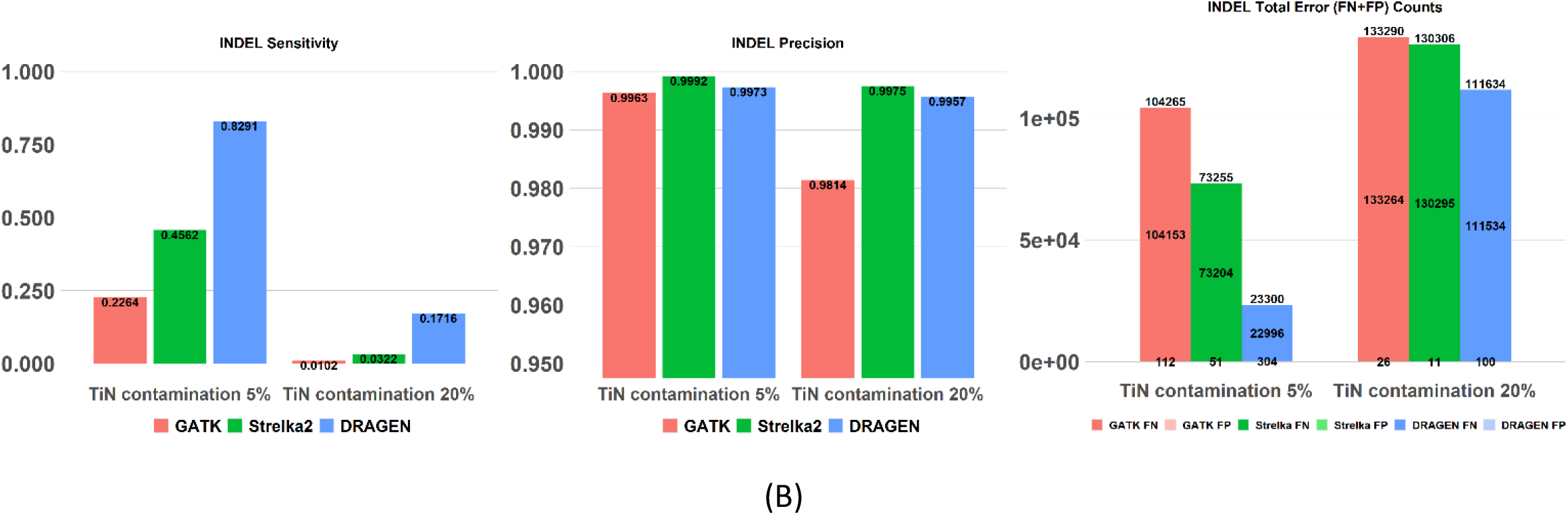
Admixture datasets with 40% tumor purity and 5% or 20% TiN contamination, comparing DRAGEN (liquid tumor mode), Mutect2, and Strelka2.

The overall summary of DRAGEN accuracy is that DRAGEN provides precision gains on the tumor cell lines datasets across the board over both other callers and for both SNV and indels. The SNV sensitivity gains are data set dependent: DRAGEN provides noticeable gains on the two NYGC cell lines that have more than 98% of truth variants > 5%, over both other callers. For the high purity SEQC2 cell line, DRAGEN is similar in SNV sensitivity to Strelka2, and both tools show gains over Mutect2. DRAGEN provides indel sensitivity gains across the board, over both other callers and across all datasets.

The germline admixture results show that DRAGEN is robust to support both low tumor purity levels and TiN contamination.

### 4.3 Run times comparison

We compared the somatic variant calling runtime of DRAGEN and Strelka2 as shown in Figure 10. The runtime for GATK was beyond 24 hours and as such was not included in the comparison. All run times were measured on the same hardware, but only DRAGEN made use of the FPGA chip that was available. The DRAGEN runtime is for a run performing map-align followed by variant calling, whereas the Strelka2 runtime is for a map-align run in DRAGEN followed by a Strelka2 variant calling run from the resulting BAM file. The runtime was measured on a DRAGEN server with two Intel Xeon Gold 6126 CPUs (total 24 cores, 2.66 GHz), a Xilinx Virtex-7 980t FPGA board, and 384 GB memory. The results show that when DRAGEN is run from FASTQ and streams the mapped reads directly into the variant caller, a T/N 100x/40x completes in about one hour. Adding the BAM output option does increase the run time but still completes under two hours.

**Figure 10.**
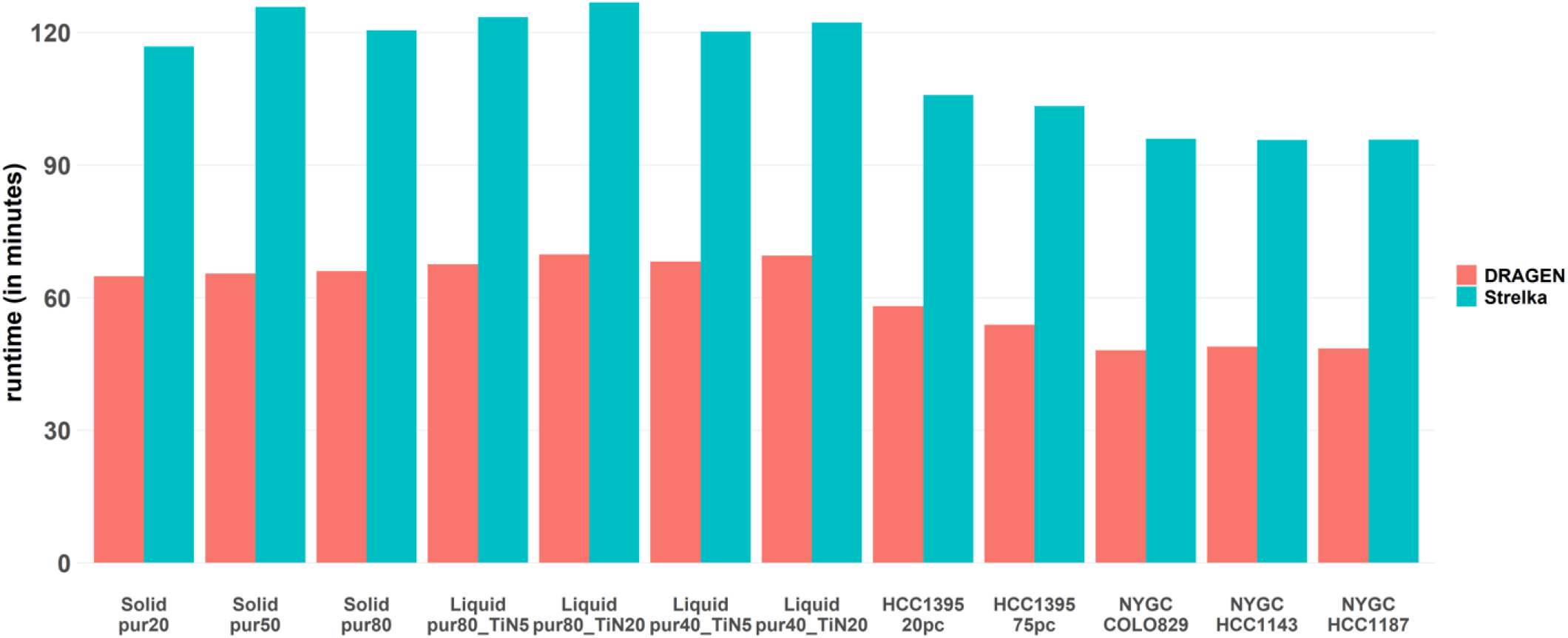
End-to-end runtime comparison. The DRAGEN runtime is for a run performing map-align followed by variant calling, whereas the Strelka2 runtime is for a map-align run in DRAGEN followed by a Strelka2 variant calling run from the resulting BAM file. The runtime was measured on a DRAGEN server with two Intel Xeon Gold 6126 CPUs (total 24 cores, 2.66 GHz), a Xilinx Virtex-7 980t FPGA board, and 384 GB memory.

## 5 Conclusion

Our results demonstrate that DRAGEN secondary analysis outperforms other state-of-the-art solutions in terms of both accuracy and speed. Although not shown here, we have also verified that DRAGEN is robust against variations in coverage, sequencing platform, and sample preparation chemistry.

The DRAGEN somatic pipeline provides seamless processing of any sequencing data type, from small deeply sequenced oncopanels, with or without UMI, to whole-genome sequencing, and allowing either paired T/N samples or tumor-only analysis. Together with FPGA-based hardware acceleration of several core algorithms in the aligner and variant caller, this results in the fastest available pipeline for processing any somatic sequencing data with top accuracy. The speed of the pipeline enables reliable whole genome analysis that can be scaled to large numbers of samples, and which we anticipate is an invaluable tool for oncology applications. DRAGEN can be run on a local server, on instrument, or in the cloud via https://basespace.illumina.com or on https://www.illumina.com/products/by-type/informatics-products/connected-analytics.html.

## 6 Somatic SNV callers

For all of the datasets presented in Table 1, we ran DRAGEN (4.0.3), Strelka2 (2.9.9), and Mutect2 (GATK4.1.3) using the reference genome hg38-alt-masked-v2 [27] and measured precision and recall of somatic-variant calls using RTG vcfeval (3.9.1). For the admixture datasets with tumor-in-normal contamination, DRAGEN and Strelka2 were run in liquid tumor mode.

DRAGEN command line:

~~~
dragen \
--output-directory $OUTPUT \
--output-file-prefix $PREFIX \
--tumor-fastq-list $TUMOR_FASTQ_LIST \
--tumor-fastq-list-sample-id $TUMOR_FASTQ_LIST_SAMPLE_ID \
--fastq-list $FASTQ_LIST \
--fastq-list-sample-id $FASTQ_LIST_SAMPLE_ID \
--ref-dir $DRAGEN_HASH_TABLE
--enable-map-align=true \
--enable-sort=true \
--enable-duplicate-marking=true \
--enable-variant-caller=true
~~~

Added to run the admixture datasets with tumor-in-normal contamination:

--vc-enable-liquid-tumor-mode true\

Both Strelka2 and GATK4/Mutect2 were run on DRAGEN 4.0.3 BAM inputs.

GATK4/Mutect2 command line

gatk Mutect2 -R ref.fasta -I tumor.bam -I normal.bam -O unfiltered.vcf followed by

gatk FilterMutectCalls -R ref.fasta -V unfiltered.vcf -O filtered.vcf

Strelka2 command line

~~~
# configuration
${STRELKA_INSTALL_PATH}/bin/configureStrelkaSomaticWorkflow.py
--normalBam normal.bam \
--tumorBam tumor.bam \
--referenceFasta hg38.fa \
--runDir demo_somatic
# execution on a single local machine with N parallel jobs demo_somatic/runWorkflow.py -m local -j 20
~~~

## 7 Data availability

NYGC COLO829, HCC1143 and HCC1185 data access is described in [17]. The raw data is available on [dbGAP](https://www.ncbi.nlm.nih.gov/gap/?term=phs001839) [18]. The somatic variant files, obtained from the high-coverage data and the somatic variant files obtained from down-sampled 80X/40X coverage, are accessible in [17].

HCC1395 and HCC1395BL data access is described in [7] and [28]. All raw data (FASTQ files) are available on NCBI’s SRA database (SRP162370). The call set for somatic mutations in HCC1395 is available on NCBI’s ftp site [21]. For precision measurement, the normal/normal dataset was built from 1) merging samples SRR7890859 and SRR7890857 together and then down-sample to 85x with samtools to generate the “tumor” sample of the N/N pair. SRR7890858 (45x) was used as the “normal” sample of the N/N pair.

Admixture datasets (mixture of NA12877 and NA12878 readsets) and their associated truth set were upload to a public S3 bucket [24].

For Research Use Only. Not for use in diagnostic procedures.

## References

[1] Cibulskis, K., Lawrence, M.S., Carter, S.L. et al.: Sensitive detection of somatic point mutations in impure and heterogeneous cancer samples. Nat. Biotechnol. 31, 213–219 (2013) doi: 10.1038/nbt.2514.

[2] Kim, S., Scheffler, K., Halpern, A.L. et al.: Strelka2: fast and accurate calling of germline and somatic variants. Nat. Methods 15, 591–594 (2018) doi:10.1038/s41592-018-0051-x.

[3] Priestley, P., Baber, J., Lolkema, M.P. et al.: Pan-cancer whole-genome analyses of metastatic solid tumours. Nature (2019) doi:10.1038/s41586-019-1689-y.

[4] GDC Data Portal with TCGA statistics: https://portal.gdc.cancer.gov/

[5] https://www.illumina.com/content/dam/illumina/gcs/assembled-assets/marketing-literature/trusight-oncology-500-data-sheet-m-gl-00173/trusight-oncology-500-and-ht-data-sheet-m-gl-00173.pdf

[6] https://emea.illumina.com/content/dam/illumina/gcs/assembled-assets/marketing-literature/trusight-oncology-500-ctdna-data-sheet-m-gl-00843/trusight-oncology-500-ctdna-data-sheet-m-gl-00843.pdf

[7] Toward best practice in cancer mutation detection with whole-genome and whole-exome sequencing, https://www.ncbi.nlm.nih.gov/pmc/articles/PMC8506910/

[8] Accuracy Improvements in Germline Small Variant Calling with the DRAGEN Platform https://science-docs.illumina.com/documents/Informatics/dragen-v3-accuracy-appnote-html-970-2019-006/Content/Source/Informatics/Dragen/dragen-v3-accuracy-appnote-970-2019-006/dragen-v3-accuracy-appnote-970-2019-006.html

[9] Cooke, D.P., Wedge, D.C., Lunter, G.: A unified haplotype-based method for accurate and comprehensive variant calling. Nat. Biotechnol. 39, 885–892 (2021) doi: 10.1038/s41587-021-00861-3.

[10] Krishnamachari, K., Lu, D., Swift-Scott, A. et al.: Accurate somatic variant detection using weakly supervised deep learning. Nat Commun 13, 4248 (2022). doi:10.1038/s41467-022-31765-8.

[11] Lai, Z., Markovets, A., Ahdesmaki M. et al.: VarDict: a novel and versatile variant caller for next-generation sequencing in cancer research. Nucleic Acids Res. 44, 11 (2016). doi:10.1093/nar/gkw227.

[12] Narzisi, G., Corvelo, A., Arora, K. et al.: Genome-wide somatic variant calling using localized colored de Bruijn graphs. Commun. Biol. 1, 20 (2018) doi:10.1038/s42003-018-0023-9.

[13] Poplin, R.,Ruano-Rubio, V., Depristo, M. et al.: Scaling accurate genetic variant discovery to tens of thousands of samples. bioRxiv (2017) doi: 10.1101/201178.

[14] Sahraeian, S.M.E., Liu, R., Lau, B. et al.: Deep convolutional neural networks for accurate somatic mutation detection. Nat Commun 10, 1041 (2019). doi:10.1038/s41467-019-09027-x.

[15] Taylor-Weiner, A., Stewart, C., Giordano, T. et al.: DeTiN: overcoming tumor-in-normal contamination. Nat. Methods 15, 7 531–534 (2018) doi:10.1038/s41592-018-0036-9.

[16] Arora, K., Shah, M., Johnson, M. et al.: Deep whole-genome sequencing of 3 cancer cell lines on 2 sequencing platforms. Sci Rep 9, 19123 (2019) doi:10.1038/s41598-019-55636-3. https://www.nature.com/articles/s41598-019-55636-3

[17] https://www.nygenome.org/bioinformatics/3-cancer-cell-lines-on-2-sequencers/

[18] [dbGAP](https://www.ncbi.nlm.nih.gov/gap/?term=phs001839)

[19] Mercer, T.R., Xu, J., Mason, C.E. et al.: The Sequencing Quality Control 2 study: establishing community standards for sequencing in precision medicine. Genome Biol 22, 306 (2021). doi:10.1186/s13059-021-02528-3.

[20] Establishing reference samples for detection of somatic mutations and germline variants with NGS technologies; https://www.biorxiv.org/content/10.1101/625624v3.full.pdf

[21] http://ftp-trace.ncbi.nlm.nih.gov/ReferenceSamples/seqc/Somatic_Mutation_WG/release/v1.2

[22] Dausset, J., Cann, H., Cohen, D. et al.: Centre d’etude du polymorphisme humain (CEPH): collaborative genetic mapping of the human genome. Genomics 6 3: 575–577 (1990). doi:10.1016/0888-7543(90)90491-c

[23] Cleary, J.G., Braithwaite, R., Gaastra, K. et al.: Comparing Variant Call Files for Performance Benchmarking of Next-Generation Sequencing Variant Calling Pipelines. bioRxiv (2015) doi: 10.1101/023754.

[24] S3 bucket with admixture datasets. To access from AWS cli: aws s3 ls -–no-sign-request s3://dragen-wgs-tn-somatic-benchmarking-datasets/

[25] NCTR Indel Calling from Oncopanel Sequencing Challenge Phase 2: https://precision.fda.gov/challenges/22/results

[26] DRAGEN High Accuracy Indel Calling Wins PrecisionFDA NCTR Indel Calling from Oncopanel Sequencing Data Challenge https://www.illumina.com/science/genomics-research/articles/precisionfda-indel.html

[27] Demystifying the Versions of GRCh38/hg38 Reference Genomes, How They are Used in DRAGEN and Their Impact on Accuracy https://www.illumina.com/science/genomics-research/articles/dragen-demystifying-reference-genomes.html

[28] Establishing reference samples for detection of somatic 2 mutations and germline variants with NGS technologies; https://www.biorxiv.org/content/10.1101/625624v1.full.pdf

